# Electron Event Representation (EER) data enables efficient cryoEM file storage with full preservation of spatial and temporal resolution

**DOI:** 10.1101/2020.04.28.066795

**Authors:** Hui Guo, Erik Franken, Yuchen Deng, Samir Benlekbir, Garbi Singla Lezcano, Bart Janssen, Lingbo Yu, Zev A. Ripstein, Yong Zi Tan, John L. Rubinstein

## Abstract

Direct detector device (DDD) cameras have revolutionized electron cryomicroscopy (cryoEM) with their high detective quantum efficiency (DQE) and output of movie data. A high ratio of camera frame rate (frames/sec) to camera exposure rate (electrons/pixel/sec) allows electron counting, which further improves DQE and enables recording of super-resolution information. Movie output also allows for computational correction of specimen movement and compensation for radiation damage. However, these movies come at the cost of producing large volumes of data. It is common practice to sum groups of successive camera frames to reduce the final frame rate, and therefore file size, to one suitable for storage and image processing. This reduction in the camera’s temporal resolution requires decisions to be made during data acquisition that may result in the loss of information that could have been advantageous during image analysis. Here we present experimental analysis of a new Electron Event Representation (EER) data format for electron counting DDD movies, which is enabled by new hardware developed by Thermo Fisher Scientific for their Falcon DDD cameras. This format enables recording of DDD movies at the raw camera frame rate without sacrificing either spatial or temporal resolution. Experimental data demonstrate that the method retains super-resolution information and allows correction of specimen movement at the physical frame rate of the camera while maintaining manageable file sizes. The EER format will enable the development of new methods that can utilize the full spatial and temporal resolution of DDD cameras.

## Introduction

Complementary metal oxide semiconductor (CMOS) direct detector device (DDD) cameras for cryoEM provide improved detective quantum efficiency (DQE) compared to other detectors (McMullan *et al.*, 2016). Furthermore, these cameras can record movies of the specimen during irradiation. Movies are output from the detector as raw ‘camera frames’ (Fig. 1A), with successive frames summed to produce ‘exposure fractions’ that are saved for image processing (Fig. 1B). Movie output has three advantages (Li *et al.*, 2013; Campbell *et al.*, 2012). First, it facilitates further improvement of DQE through the implementation of electron counting, where an algorithm is used to detect, localize, and normalize the signal from each electron in individual camera frames. Second, it allows super-resolution imaging by recording the positions of electrons with an accuracy finer than size of the sensor’s physical pixels. Finally, DDD movies makes it possible to account for radiation damage to the specimen and correct the beam-induced specimen motion and microscope stage drift that occur during imaging. DQE is improved by electron counting because the signal contributed to the image by each electron varies stochastically (McMullan, Faruqi *et al.*, 2009) and consequently counting electrons normalizes this signal (Li *et al.*, 2013). For electron counting, the exposure per frame is limited to one electron for every ~40 to 100 pixels. This low density of electrons per frame allows individual electrons to be detected with a low probability of two electrons impinging on the same region during the recording of the frame, which would lead to undercounting electrons in a phenomenon known as ‘coincidence loss’. Each electron deposits energy into multiple pixels upon hitting the sensor, and consequently the center of the impact event can be localized to a specific region of a pixel in order to allow super-resolution imaging (Li *et al.*, 2013). Recording super-resolution information also improves the DQE of the camera within the physical Nyquist frequency by reducing noise aliasing (McMullan, Chen *et al.*, 2009).

**Figure 1.**
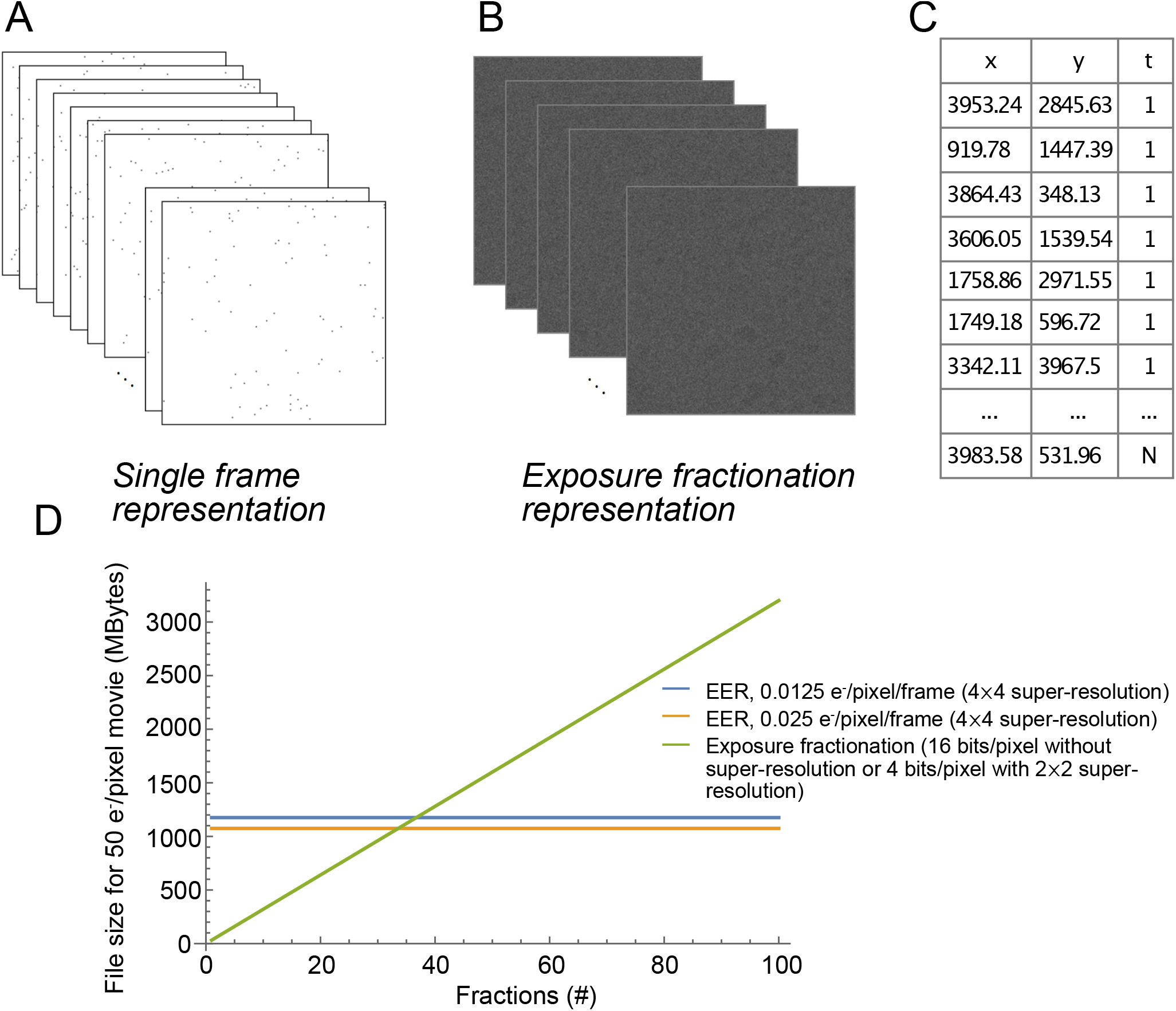
The EER file format. **A,** Direct detector device (DDD) cameras operating in counting mode record the impact positions of electrons on the sensor at the frame rate of the camera. **B,** Conventionally, groups of movie frames are averaged to fractionate the exposure, reducing the size of movie files from DDD cameras. This exposure fractionation requires decisions to be made by the experimentalist about the temporal resolution to be preserved in order to avoid loss of information from specimen movement during imaging. **C,** The electron event representation (EER) file format uses efficient data encoding, marking the position and time (in raw frame number) for each electron. **D,** Example data sizes under typical conditions. All reported data sizes assume a total exposure on the specimen of 50 e^−^/Å^2^, a pixel size of 1 Å, frame size 4096×4096 pixels, and neglect any loss of electrons between specimen exposure and detection with the camera. *Green curve*: data size for 16 bits/pixel or (equivalently) 4 bits/pixel with 2×2 super-resolution. *Blue* and *orange* curves: EER file sizes with 4×4 super-resolution at an exposure rate of 0.0125 e^−^/Å^2^/frame and 0.025 e^−^/Å^2^/frame, respectively. The EER file size depends only on the total electron exposure and exposure rate of the camera, while the file size for conventional movies depends on the number of fractions recorded. EER thus preserves the full temporal resolution of the electron detection events and requires a smaller file size for many practical fractionation conditions.

Beam-induced motion and specimen drift, which blur images of ice-embedded protein complexes in integrated exposures, can limit attainable resolution by cryoEM. Numerous schemes have now been implemented to correct this motion (Ripstein & Rubinstein, 2016). The earliest approaches treated the image on the entire area of the detector as moving in unison (Brilot *et al.*, 2012; Li *et al.*, 2013). Later approaches divide the detector into patches (Zheng *et al.*, 2017) or work on individual particle images, using either the shift-dependent average of exposure fractions (Rubinstein & Brubaker, 2015) or a projection of a 3D map (Zivanov *et al.*, 2019) to guide alignment. Finally, radiation damage to specimens means that the early part of each exposure contains more high-resolution information than the later part (Baker *et al.*, 2010), and this loss of information can be accounted for when averaging exposure fractions (Rubinstein & Brubaker, 2015; Feng *et al.*, 2017; Grant & Grigorieff, 2015) or during 3D reconstruction (Scheres, 2014; Zivanov *et al.*, 2019).

The smallest possible exposure fraction from a camera is a single camera frame, with current hardware frame rates for ~4k×4k pixel sensors of between 40 and 1500 frames/sec. Consequently, camera movie modes have the potential to produce enormous volumes of data. For example, a 4096×4096 pixel sensor with a readout rate of 400 frames/sec and with pixel values stored as 4 bits of information would produce 3.125 GiB of information each second. Movies must be recorded over multiple seconds for electron counting with an appropriate total electron exposure and magnification for a 2 to 3 Å resolution reconstructions of a biological specimen (Ripstein & Rubinstein, 2016). Therefore, while DDDs have revolutionized cryoEM and structural biology as a whole, they have placed great demands on current computational data storage infrastructure. Because storing the entirety of these movies is not practical, experimentalists must make decisions not just about magnification (Å/pixel), total electron exposure on the sample (e^−^/Å^2^), and camera exposure rate (e^−^/pixel/second), but also about how to best fractionate exposures by summing successive frames after electron counting. If exposures are fractionated too finely, file sizes are excessively large. If exposures are fractionated too coarsely, significant motion can occur within one fraction, compromising the resolution of 3D structures that can be calculated from the data. These decisions are made at the time of data collection and the microscopist runs the risk of realizing during analysis that their data acquisition strategy was not optimal.

In this paper we describe Electron Event Representation (EER), an image recording strategy developed at Thermo Fisher Scientific for their Falcon cameras. We show that storing EER data removes the need to decide on an exposure fractionation strategy during imaging, enabling optimal correction of specimen motion. In addition, we demonstrate that EER files record super-resolution information in images, allowing 3D reconstruction beyond the Nyquist frequency.

## Results

### Theoretical basis for EER

Conventional representations of cryoEM movies store pixel intensities for each exposure fraction. In contrast, in EER each electron detection event is recorded as a tuple of position and time (*x*,*y*,*time*), indicating where and when the electron was detected on the sensor (Fig. 1C). As discussed earlier, due to the need to avoid coincidence loss during electron counting, the number of detected electrons in a single camera frame must be ~40 to 100 times smaller than the number of pixels in the frame. This inherent sparsity may be exploited for efficient encoding of pixel locations for the detected electrons. Assuming that in a single electron counted camera frame, each pixel is either not hit (value 0) or hit (value 1) by an electron, the stream of camera frame pixels can be modeled as a Bernoulli process with the probability *p* of an individual pixel being hit by an electron given by

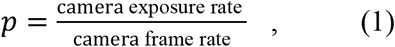

where the camera exposure rate has dimensions e^−^/pixel/sec and the frame rate has dimensions frames/sec. The Shannon entropy (Shannon, 1948), *H*, of this Bernoulli process is

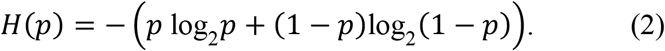

This Shannon entropy gives a lower bound on the number of bits per pixel needed to encode all events in a counted frame. Reaching this lower bound requires that the statistical model matches the statistics of the data and that an optimal data compression scheme is used. A value of *p ≠ 0.5* leads to *H(p)<1* and indicates that the camera frames can be compressed further. Recording electron locations on the sensor with super-resolution accuracy by subdivision of physical pixels into *u×u* sub-pixels requires 2 log_2_(*u*) additional bits per electron. Consequently, the size in bytes, *D*, for an optimally compressed EER movie frame is given by

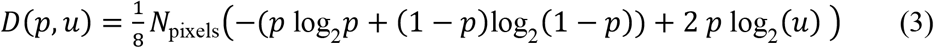

where *N*_pixels_ is the number of physical pixels in the sensor. For example, on a sensor with 4096×4096 pixels running at a frame rate of 240 frames/sec, a camera exposure rate of 3 e^−^/pixel/sec gives *p* = 3/240 = 0.0125. When each pixel is sub-divided into 4×4 sub-pixels (*u*=4), an optimally compressed EER movie requires 301 kB/frame. Without recording super-resolution location information (*u*=1) the same EER movie would require 199 kB/frame. The expected total size *S*_*opt*_ of an optimally compressed EER movie in bytes, neglecting any file header information, is therefore given by

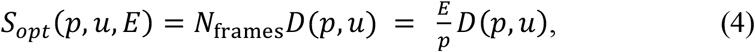

where *E* is the total electron exposure in the movie in e^−^/pixel and *N*_frames_ is the number of camera frames recorded.

The EER format implemented for Falcon cameras uses run-length encoding (RLE) to reduce data size. For each camera frame the pixel distances between detected electrons, in the scanline order in which they are stored in memory, are encoded with a constant word length, *b*_*RLE*_. In the current algorithm, *b*_*RLE*_ was set at 7 bits. The maximum value, *m*, for the given number of bits (i.e. 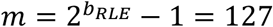 for *b*_*RLE*_*=*7 bits) is used to indicate that there was no electron detected after this maximum number of *m* pixels. This scheme does not achieve the optimal data compression and file size described in equation 4, but has the advantage of straightforward image encoding and decoding. The approximate total file size with RLE compression, *S*_*RLE*_, is given by the product of total electron exposure *E*, number of pixels *N*_pixels_, and the number of bits per electron *b*_*RLE*_ + 2 log_2_(*u*), but with a correction to account for the extra bits needed to represent the situation where no electron was detected after *m* pixels:

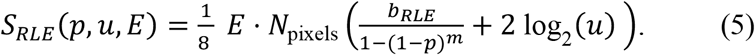

The optimal choice for *b*_*RLE*_ to minimize file size depends on *p*. The use of 7 bits enables small file sizes when typical exposure rates for electron counting are used. The EER format implemented for Falcon cameras uses *u*=4, meaning physical pixels are divided into 4×4 sub-pixels.

Figure 1D shows typical EER file sizes (50 e^−^/pixel total exposure with 1 Å/pixel) compared to standard image formats, such as MRC image stack files (Cheng *et al.*, 2015). In contrast to the EER files, the MRC files described in the figure have reduced temporal resolution due to averaging of successive frames. Where the example MRC files preserve super-resolution information they use 2×2, rather than 4×4, sub-pixels. When more than ~35 exposure fractions are recorded, EER files are smaller than 16-bit MRC files or 4-bit MRC files with 2×2 super-resolution information. The intersection of the EER curve with the conventional fractionation approach curve will occur at a larger number of exposure fractions if a compressed image format is used (e.g. LZW-TIFF). However, the amount of image compression that can be achieved depends strongly on image content and consequently it is difficult to compare these methods analytically. In principle, RLE compression could be applied to conventional movies saved with each exposure fraction consisting of a single super-resolution camera frame. However, the real-time output of EER data from the camera avoids saving extremely large uncompressed intermediate files even temporarily, which would make workflows prohibitively complicated. Lossy compression approaches have also been shown to reduce file sizes when complete preservation of information is not required (Eng *et al.*, 2019). Consequently, conventional files that are smaller than the EER format can be produced, but doing so requires sacrificing temporal or spatial resolution.

### Super-resolution imaging

Modern DDD cameras such as the Gatan K2 or K3, Direct Electron DE-16 or DE-64, and Thermo Fisher Scientific Falcon 3EC or 4 localize electrons with sub-pixel accuracy using a centroiding procedure before electron positions are recorded. As described above, this super-resolution information is preserved in the EER format by sub-dividing each physical pixel into *u×u* sub-pixels. Because the Nyquist resolution of a camera is given by two times the edge length of a pixel, sub-division of physical pixels by a factor of *u* extends the Nyquist resolution by 1/*u*. Even without sub-pixel localization of electrons, images retain information beyond the Nyquist frequency because the corners of Fourier transforms encode spatial frequencies that are finer than the Nyquist frequency in the *x* or *y* direction of the image. (Fig. 2A).

**Figure 2.**
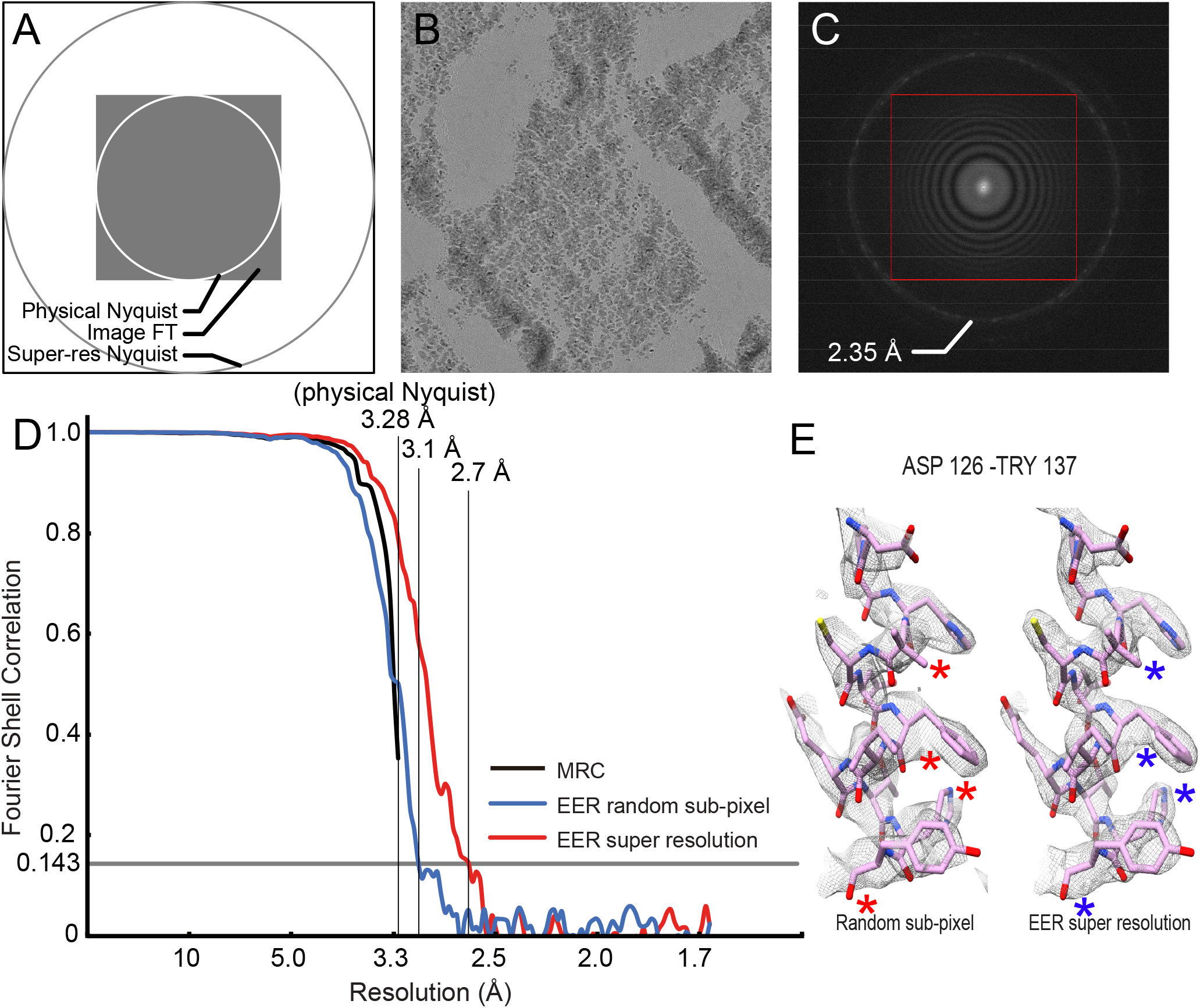
Super-resolution 3D reconstruction with EER files. **A,** Illustration of the physical Nyquist frequency, information in square Fourier transforms beyond the physical Nyquist, and the new Nyquist frequency from 2×2 supersampling of physical pixels. **B,** Image of a cross-grating with polycrystalline gold recorded as an EER file. **C,** Fourier transform of the image from part A, showing information present outside of the Fourier transform of the image’s physical pixels (*red box*) and beyond the physical Nyquist frequency (*red circle*). **D,** FSC curves from maps with a physical Nyquist resolution of 3.28 Å: standard images (*black curve*), 2×2 supersampled with random sub-pixel electron placement (*blue curve*), and 2×2 supersampled with sub-pixel electron placement from the EER file (*red curve*). **E,** Part of an α helix from a 3D map at 3.1 Å resolution (FSC=0.143) from random sub-pixel information (*left*) and at 2.7 Å resolution (*right*) with super-resolution information from EER data. Asterisks (*) indicate features that are better resolved on the *right* than on the *left*.

We investigated the ability of a Titan Krios electron microscope with a Falcon camera and EER capability to record information beyond the physical Nyquist frequency of the camera sensor. Images of a standard cross-grating with polycrystalline gold were recorded with a physical pixel size of 1.71 Å (Fig. 2B). The Fourier transform of the image shows diffraction peaks that correspond to 2.35 Å, or 1.46× the Nyquist resolution of 3.42 Å (Fig. 2C, *red circle*). Therefore, it is evident that the electron counting algorithm combined with the EER data format enables recording of information beyond the physical Nyquist limit of the camera.

To test whether the super-resolution capability of EER files could be applied to biological specimens, we imaged human light-chain apoferritin particles with a calibrated physical pixel size of 1.64 Å and a physical pixel Nyquist resolution of 3.28 Å. Movies were recorded as EER data with a total exposure of ~42 e^−^/Å^2^ on the specimen and a camera exposure rate 0.63 e^−^/pixel/sec. These movies were then converted to 30 MRC format exposure fractions. 3D reconstruction from 118,766 particle images extracted from 157 movies with a conventional refinement work-flow gave a 3D resolution by Fourier shell correlation of 3.3 Å (Fig. 2D, *black curve*). It should be noted that 3D reconstructions with resolutions close to the Nyquist frequency can suffer from artefacts that limit the ability to resolve their highest-resolution features. Next, the same EER files were converted to movies with 30 fractions but with a pixel size of 0.82 Å (Nyquist resolution 1.64 Å). Electrons were placed on pixel grid that is 4×4 supersampled from the camera’s physical pixel grid. Sub-pixel positions were either chosen randomly or using the EER information. Subsequently, the image were Fourier cropped to give an effective 2×2 supersampling of the physical pixel grid. 3D reconstruction from these images following the same workflow used with the conventional image files gave 3D maps with resolutions of 3.1 Å for the random sub-pixel placement (Fig. 2D, *blue curve*) and 2.7 Å for placement with information from EER (Fig. 2D, *red curve*). The resolution from the randomized sub-pixel information, 3.1 Å, is notable because it goes beyond the physical Nyquist resolution of 3.28 Å. This effect is due to information past the Nyquist resolution found in the corners of the Fourier transform of the image (Fig. 2A), although improved motion correction in the supersampled images may also improve the map. The resolution from the reconstruction that used sub-pixel information from the EER file was 2.7 Å, 18 bins in Fourier space beyond the physical Nyquist resolution and 13 bins in Fourier space beyond the randomized sub-pixel control. Numerous features in the maps indicate improved resolution where EER sub-pixel information was used (Fig. 2E, *right*, blue asterisks) compared to where random information was used (Fig. 2E, *left*, red asterisks).

### Intra-fraction motion correction enabled by EER imaging

The ability to fractionate exposures up to the physical frame rate of the camera, without needing to store the data as high frame rate movies, provides the possibility of improved measurement and correction of beam induced motion. However, estimating motion from extremely large numbers of fractions can be problematic for the current generation of motion measurement algorithms (Rubinstein & Brubaker, 2015; Zivanov *et al.*, 2019; Zheng *et al.*, 2017). Alternatively, motion can be measured from a smaller number of fractions but the trajectory subsequently interpolated or extrapolated to the raw camera frames.

Using the implementation of the *alignparts_lmbfgs* algorithm (Rubinstein & Brubaker, 2015) in *cryoSPARC* (Punjani *et al.*, 2017), we measured the motion trajectory of 291,408 single particle images of apoferritin. These trajectories were measured in EER movies that had been divided into 30 exposure fractions, where each exposure fraction was comprised of 77 camera frames. Images were recorded with a calibrated physical pixel size of 1.06 Å but supersampled 1.5×1.5 to super-resolution pixels of 0.7067 Å with information from the EER data. To mimic conventional movie processing, the motion measured from the 30 exposure fractions was applied uniformly to all of the frames within each fraction (Fig. 3A, *yellow line*). Exposure weighting, as proposed previously (Baker *et al.*, 2010), was performed as described in the *alignparts_lmbfgs* algorithm (Rubinstein & Brubaker, 2015) but using resolution-dependent optimal exposures that were measured subsequently (Grant & Grigorieff, 2015). This strategy is equivalent to the exposure weighting done with *Motioncor2* (Zheng *et al.*, 2017), *Unblur* (Grant & Grigorieff, 2015), and *cryoSPARC* (Punjani *et al.*, 2017). To assess the benefit of increased time-resolution in the applied motion trajectories, 3^rd^ order B-spline interpolation was used to assign the position of each particle in each camera frame (Fig. 3A, *blue line*). Three-dimensional reconstruction using just the measured motion from the 30 exposure fractions without interpolation produced a map at 2.10 Å resolution (Fig. 3B, *black curve*). In contrast, applying interpolated motion at the physical frame rate prior to averaging gave a map at 2.07 Å, which is an improvement of two bins in Fourier space (Fig. 3B, *red curve*). Beam-induced motion in the early frames of a movie is thought to be one of the primary limits to resolution in cryoEM at present (Henderson, 2018). This modest improvement in resolution from interpolated application of the measured motion suggests that the motion estimates from the fractionated movie are not sufficiently accurate to allow improved resolution.

**Figure 3.**
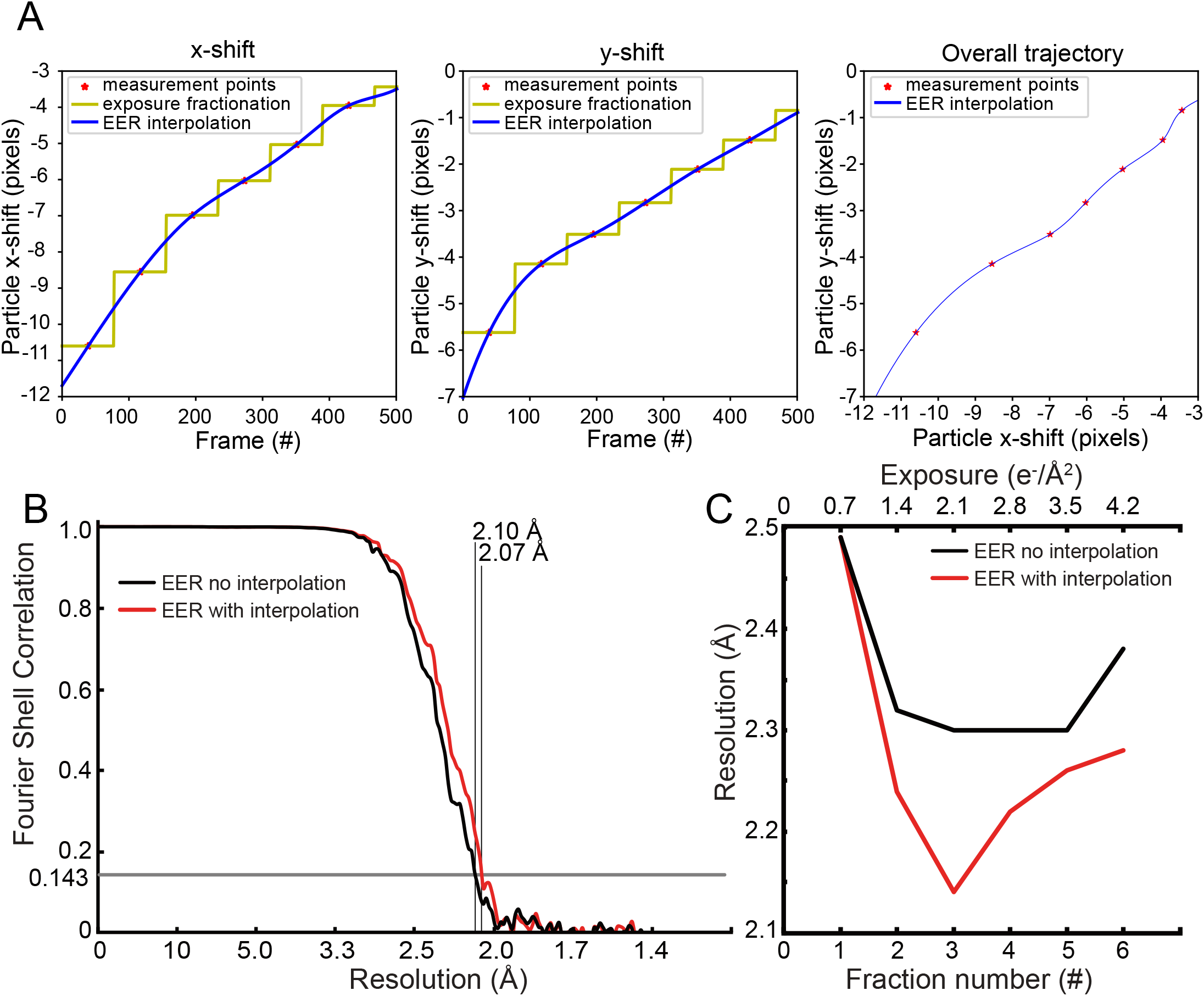
Improved correction of beam-induced motion with EER files. **A,** Example of individual particle trajectories measured from 30 exposure fractions and interpolated to the physical frame rate of the camera. The *yellow* line represents the applied motion without the B-spline interpolation enabled by the EER method while the *blue* line represents the interpolated trajectory enabled by EER. **B,** Fourier shell correlation curve for 3D reconstructions without (*black curve*; 2.10 Å resolution at FSC=0.143) and with (*red curve*; 2.07 Å resolution at FSC=0.143) interpolated motion applied at the camera frame rate. **C,** Comparison of resolution for 3D maps (FSC=0.143) calculated from different exposure fractions, each corresponding to 0.7 e^−^/Å^2^, without (*black curve*) and with (*red curve*) interpolated motion applied to the camera frames.

In contrast to the overall map resolution, the resolutions of 3D maps calculated from individual exposure fractions improved markedly when motion trajectories were interpolated and applied directly to camera frames. Movies, with each fraction consisting of 77 frames with 1.4 e^−^/Å^2^/fraction, were fractionated further to averages of 38 frames, corresponding to 0.7 e^−^/Å^2^/fraction. 3D maps were calculated separately from the first six of these new fractions, with or without the application of the motion to the individual camera frames in each fraction. During this 3D reconstruction the orientations of particle images were not changed from those measured from the exposure-weighted average of fractions. The resolutions of the resulting maps are shown in Fig. 3C. Remarkably, the resolutions of these maps are only 0.07 to 0.4 Å worse than the resolutions of the maps calculated from the exposure-weighted average of all frames from the movies. This result indicates that, while information from the entire exposure may guide alignment of particle images to a 3D reference, the high-resolution features in maps can be reconstructed from just the earliest part of the exposure. While the first fraction is no better with the interpolated motion than with the non-interpolated motion, the subsequent fractions show a marked improvement in resolution. Consequently, it appears that the estimated motion is not correct during the earliest part of the exposure where the specimen moves the most and with the least predicable direction. However, later in the exposure the estimated motion is sufficiently accurate to allow improved map resolution when the trajectory is interpolated and applied directly to the camera frames.

## Discussion

Processing of EER images in this work required an intermediate image processing step of converting EER data into a movie format that could be used by *cryoSPARC* (Punjani *et al.*, 2017) and *Relion* (Scheres, 2012), the software packages we employed for image analysis. However, information about the EER file format has already been shared with the development teams for these software packages and the capability to directly read EER has been implemented in both packages. The file format specification is also available to other software developers.

DDDs have previously allowed extraction of information beyond the physical Nyquist frequency of the camera for images of 2D crystals (Chiu *et al.*, 2015) and single particles (Feathers *et al.*, 2019), with other algorithms proposed to explore this approach further (Chen, 2018). When subdividing each physical pixel into 4×4 sub-pixels, the EER format allows preservation of super-resolution information with an additional 4 bits required for each electron detected, which increases file sizes by a maximum of 57%. In contrast, conventional representations of a super-resolution image with each physical pixel divided into 2×2 sub-pixels causes a 400% increase in file size relative to the non-super-resolution image. Dividing the physical pixel into 4×4 sub-pixels, as done in the EER format, would increase the file size by 1600%. Acquiring images at lower magnification provides more particles per image and decreases time spent preparing for the exposure. However, super-resolution imaging does not provide a dramatically faster route to high-resolution cryoEM data collection. Decreasing the microscope magnification requires keeping the camera exposure rate (e^−^/pixel/sec) constant to allow for electron counting and requires more time to obtain the same total specimen exposure (e^−^/Å^2^). Nonetheless, the preservation of super-resolution information decreases the importance of the magnification chosen when data collection is initiated. Further, a lower magnification increases the field of view in images, which can facilitate measurement of specimen tilt and the microscope contrast transfer function. A larger field of view may also improve modelling of beam induced motion, which typically utilizes information from movement of adjacent particles (Scheres, 2014; Rubinstein & Brubaker, 2015). The increased field of view can also be advantageous for electron tomography of larger objects.

The calculation of 3D maps from different exposure fractions described in Fig. 3C shows that it is possible to obtain the highest-resolution from a single exposfraction after pre-exposure of the specimen with 1.4 e^−^/Å^2^. This finding is consistent with the large body of evidence that the earliest part of the exposure, where high-resolution information should be best preserved, suffers from the most beam-induced specimen motion (Henderson, 2018). The position of this optimum indicates that smoother application of the measured particle motion from interpolation has the greatest effect near the beginning of the movie where motion is still large, while in the first 1.4 e^−^/Å^2^ of exposure inaccuracies in the measured motion prevent the smoother application from improving map resolution. This result is particularly encouraging. It suggests that new techniques that are capable of more accurate measurement of beam-induced motion could allow for extraction of high-resolution information from the earliest frames of a movie. EER data, which preserves the full temporal resolution of data acquired with DDD cameras while maintaining manageable file sizes, can allow for development of these improved beam-induced motion correction methods.

## Methods

### Specimen preparation

Human apoferritin was a gift from Ms. Taylor Sicard and Prof. Jean-Philippe Julien (The Hospital for Sick Children) and was used at 10 mg/mL. Holey gold grids with a regular array of ~2 μm holes were prepared as described previously (Marr *et al.*, 2014). Grids were subjected to 15 sec of glow discharge in air before freezing in liquid ethane with a Gatan CP3 grid freezing device. The grid freezing device chamber was at room temperature, 90 % RH, and blotting was done for 10 sec with an offset of −0.5 mm.

### Data collection

Images were acquired as described in the main text with a Titan Krios G3 electron microscope from Thermo Fisher Scientific operating at 300 kV and equipped with a Falcon 3EC camera and a prototype EER module (used for intra-fraction motion correction experiments) and later a prototype Falcon 4 camera (used for super-resolution experiments). Automatic data collection was done with the *EPU* software package. For EER intra-frame motion correction, 325 movies of human light-chain apoferritin were collected with the Falcon 3EC camera at 75,000× nominal magnification, corresponding to a calibrated pixel size of 1.06 Å. Falcon 3EC movies were recorded simultaneously in both EER format with 2312 raw frames per movie as well as 16-bit MRC format with 30 fractions per movie. The camera exposure rate and the total exposure of the specimen were 0.80 e^−^/pixel/sec and ~41 e^−^/Å^2^, respectively, with defocus ranging from 0.4 μm to 1.6 μm. Following completion of this aspect of the work, we replaced the Falcon 3EC camera with a prototype Falcon 4 camera, which increased the physical frame rate from 40 to 250 frames/sec. Consequently, for EER super-resolution data, 157 movies were collected on the same microscope but with the prototype Falcon 4 camera. A nominal magnification of 47,000× gave a calibrated pixel size of 1.64 Å. This camera did not allow for simultaneous recording of EER data and conventional movies. After collection, these EER files could be converted to standard MRC files with the desired exposure fractionation. The camera exposure rate was 4.72 e^−^/pixel/sec and the total exposure on the specimen was ~42 e^−^/Å^2^. Movies were stored in EER format with 6020 raw frames per movie. Defocus in this dataset ranged from 0.3 to 1.5 μm.

### EER image handling

The prototype EER module for Falcon 3EC camera ran custom firmware with real-time EER encoding, streaming the data to a dedicated computer running the Ubuntu 16.04 operating system. With the Falcon 4 camera, the EER files were stored with the standard Falcon 4 storage infrastructure, which normally records MRC exposure fractionation stacks. Electron detection events were stored with run-length encoding as described in the text of the manuscript. Frames were packed in a BigTIFF compliant file format with a gain reference image stored separately in an MRC file. Information about defects were encoded in the same gain reference with a value of ‘0’. EER files were decoded using a hybrid CPU/GPU implementation of the decoding algorithm. To utilize sub-pixel information optimally for both super-resolution and non-super-resolution cases, all decoded images were reconstructed on the full 4×4 supersampled image grid and subsequently Fourier-cropped to the desired resolution. For single particle cryoEM, EER files were converted to standard exposure fractionated image stacks that could be used in standard image processing pipeline. In the final correction of motion for individual particle images, the EER files were decoded with the desired supersampling (i.e. 4×4 oversampling followed by Fourier cropping), image shifts applied, and exposure-weighting performed as described previously (Rubinstein & Brubaker, 2015). Application of image shifts to data from EER files was done by placing electrons on shift-compensated positions rather than first composing an image and then applying shifts by interpolation in real space or phase changes in Fourier space. The procedure of shifting electron positions prior to image reconstruction is less expensive computationally than image interpolation, and prevents image interpolation artefacts. Efficient gain correction was performed by retrieving the gain correction coefficient from the uncorrected pixel locations for each detected electron and applying it as a weighting factor for the contribution of the electron to its shifted position. During these procedures, the individual particle motion trajectories were either smoothed with cubic spline interpolation, or not interpolated as a control, as described in the manuscript.

### Single particle cryoEM image analysis

For the Falcon 3EC dataset, 325 16-bit MRC movies were imported in *cryoSPARC* v2 (Punjani *et al.*, 2017). Movie frames were aligned with an improved implementation of *alignframes_lmbfgs* (Rubinstein & Brubaker, 2015) within *cryoSPARC* v2 and CTF parameters were estimated from the average of aligned frames with *CTFFIND4* (Rohou & Grigorieff, 2015). 335,137 particle images were selected and beam-induced motion for individual particles was corrected with an improved implementation of *alignparts_lmbfgs* (Rubinstein & Brubaker, 2015) within *cryoSPARC* v2. After two rounds of 2D classification, 291,408 particle images were selected and divided into 3 beam tilt groups. Initial homogeneous refinement was performed in *cryoSPARC* v2 without CTF refinement. The alignment information in the *cryoSPARC* .cs file was converted to *Relion* 3.0 .star file format with the *pyem* package (DOI: 10.5281/zenodo.3576630), allowing per-particle CTF and per-group beam tilt to be calculated in *Relion* 3.0. Refinement of CTF and beam-tilt parameters without alignment in *Relion* (Zivanov *et al.*, 2020) but with imposed octahedral symmetry produced a 3D reconstruction at 2.14 Å resolution. Super-resolution images of the particles with a new pixel size of 0.7067 Å were extracted with and without intra-frame motion correction as described above. Refinement of CTF and beam tilt parameters was done in *Relion* using the angles previously determined. An equivalent analysis was performed on the first six 0.70 e^−^/Å^2^/fractions of the EER movies.

For super-resolution experiments with the Falcon 4 dataset, 157 EER movies were decompressed and converted to 32-bit floating point MRC format. Movie fractions were aligned by patch-based motion correction and contrast transfer function (CTF) parameters were determined with patch CTF estimation in *cryoSPARC* v2 (Punjani *et al.*, 2017). Templates for automatic particle selection were generated by 2D classification of manually selected particles. 154,292 single particle images were selected from the aligned fractions and beam-induced motion correction for individual particles and exposure weighting was done in *cryoSPARC* v2 in the same way as described for the Falcon 3EC dataset. A subset of 118,766 particle images was selected by 2D classification and divided into four beam tilt groups. Homogeneous refinement in *cryoSPARC* v2 with imposed octahedral symmetry, per-particle defocus refinement, and higher-order aberration correction (Zivanov *et al.*, 2020), including beam tilt and trefoil aberration, yielded a map at 3.3 Å resolution. Super-resolution images of the same particles with a pixel size of 0.82 Å were extracted from EER movies with and without random sub-pixel electron placement as described above. Similar homogeneous refinement of the super-resolution particles with and without random sub-pixel electron placement yielded maps at 3.1 Å and 2.7 Å resolutions, respectively.

## Statement of contributions

E.F., B.J., and L.Y. devised the EER approach. E.F. and G.S.L. implemented the EER encoding and decoding firmware and software. JLR supervised the analysis of experimental data. JLR, HG, EF, and YD designed experiments with input from ZAR, YZT, and SB. SB prepared the apoferritin grids and imaged them with the Titan Krios microscope. HG, EF, and YD performed calculations and analysed the data. JLR, EF, and HG wrote the manuscript and prepared the figures with input from the other authors.

## Acknowledgements

We thank Xander Jansen (Thermo Fisher Scientific) for assistance with the prototype EER hardware and Falcon 4 camera in Toronto and Miloš Malínský (Thermo Fisher Scientific) for acquiring the super-resolution cross-grating EER data used in Figure 2B and 2C. This work was supported by Thermo Fisher Scientific and a Discovery Grant from the Natural Sciences and Engineering Research Council (JLR), an Ontario Graduate Scholarship (HG), a Canada Graduate Scholarship (ZAR), a postdoctoral fellowship from the Canadian Institutes of Health Research (YZT), and the Canada Research Chairs program (JLR). CryoEM data was collected at the Toronto High-Resolution High-Throughput cryoEM facility, supported by the Canada Foundation for Innovation and Ontario Research Fund. EF, YD, GSL, BJ, and LY are employees of Thermo Fisher Scientific. JLR is an advisor to Structura Biotechnology Inc.

